# First Genome-Scale Metabolic Model for Understanding HMPV-Host Interaction

**DOI:** 10.64898/2026.06.10.731436

**Authors:** Syed Mushahid Hussain, Lukas Beierle, Bernd Schmeck, Reihaneh Mostolizadeh

**Affiliations:** Mathematics and Computer Science, Philipps University of Marburg, 35032 Marburg, Germany; Institute for Lung Research, German Center for Lung Research (DZL), Universities of Giessen and Marburg Lung Center, Philipps University of Marburg, Marburg, Germany; Medicine, Pulmonary and Critical Care Medicine, University Medical Center Marburg, Philipps University of Marburg, German Center for Lung Research (DZL), Marburg, Germany; German Center for Infection Research (DZIF), the Center for Synthetic Microbiology (SYNMIKRO), Marburg, Germany, and the Institute for Lung Health (ILH), Giessen, Germany; Bioinformatics and Systems Biology, Justus Liebig University Giessen, 35392 Giessen, Germany; Biochemistry and Synthetic Metabolism, Max-Planck-Institute for Terrestrial Microbiology, AG Erb, 35043 Marburg, Germany

**Keywords:** Human Metapneumovirus (HMPV), Genome-scale metabolic model (GEM), Viral Biomass Objective Function (VBOF), Flux balance analysis (FBA), Antiviral targets, Hexosamine biosynthesis pathway (HBP)

## Abstract

**Background:** Human Metapneumovirus (HMPV) is a major contributor to acute respiratory tract infections. Currently, no approved vaccines or specific antiviral therapies are available worldwide. Genome-scale metabolic models (GEMs), when integrated with Viral Biomass Objective Functions (VBOFs), provide a robust computational framework for identifying host metabolic dependencies essential for viral replication. This approach enables systematic prioritization of potential antiviral drug targets.

**Results:** The first comprehensive VBOF for HMPV was constructed by integrating stoichiometric data from the viral genome, including structural proteins with defined copy numbers, amino acid residues per virion, envelope lipids, and glycan modifications. The reconstructed VBOF was incorporated into the human bronchial epithelial cell model, *i*HBEC^1^, to analyze metabolic changes between uninfected and infected host cells. Knockout analysis identified two selective antiviral gene targets: `PGM3` (phosphoacetylglucosamine mutase) and `GNPNAT1` (N-acetylglucosamine-6-phosphate acetyltransferase), as well as seven selective reaction targets primarily within the hexosamine biosynthesis and nucleotide sugar pathways. Knockout of `PGM3` or `GNPNAT1` completely abolished viral production while preserving complete host cell viability. Additionally, guanylate kinase (GUK1/GK1) emerged as a highly selective target for HMPV, confirming findings from Severe Acute Respiratory Syndrome Coronavirus 2 (SARS-CoV-2) studies and suggesting a conserved vulnerability across respiratory viruses.

**Conclusions:** UDP-GlcNAc, the vital end-product of the hexosamine biosynthesis pathway (HBP), is computationally predicted as a critical metabolic hub. This pathway represents the primary metabolic vulnerability of HMPV, due to the extensive glycosylation requirements of the HMPV attachment protein. The host-directed antiviral targets `PGM3` and `GNPNAT1` are high-priority candidates for experimental validation and may provide novel strategies to combat this respiratory infection.

## 1 Introduction

Human Metapneumovirus (HMPV) is an enveloped, negative-sense, single-stranded ribonucleic acid (RNA) virus classified within the *Pneumoviridae* family. It was first identified in 2001^2^. HMPV is a significant cause of acute respiratory tract infections, particularly among young children, older adults, and immunocompromised individuals^3, 4^. Clinical presentations range from mild upper respiratory symptoms to severe bronchiolitis and pneumonia. HMPV frequently co-circulates with respiratory syncytial virus Respiratory Syncytial Virus (RSV) and influenza, and these viruses exhibit comparable mortality rates^4, 5^. Mortality rates are significantly higher among infants residing in low- and lower-middle-income countries^6^. In December 2024, a seasonal outbreak of HMPV resulted in a substantial increase in hospitalizations in China.

Lessons learned from the rapid global spread of COVID-19 have heightened public awareness of the potential for novel viruses to spread rapidly^6^. The World Health Organization (WHO) and Chinese authorities have consistently stated that the emerging cases remain within the expected range for the winter season and that there is currently no indication of a pandemic^6^. Despite the considerable global disease burden posed by HMPV, no approved vaccines or specific antiviral therapies are currently available for HMPV infection.

The HMPV genome consists of approximately 13,350 kilobase(Kb). HMPV encodes nine proteins: nucleoprotein (N), phosphoprotein (P), matrix protein (M), fusion protein (F), attachment glycoprotein (G), small hydrophobic protein (SH), RNA-dependent large polymerase protein (L), and two regulatory proteins, M2-1 and M2-2^7^. The virion structure is pleomorphic, with spherical particles having a mean diameter of 209 nanometer(nm), and filamentous forms averaging 282 nm in length^7^. Similar to other paramyxoviruses, HMPV depends extensively on host cell machinery for replication. These host-dependent processes include nucleotide synthesis for genome replication, utilization of amino acid pools for protein synthesis, lipid biosynthesis for envelope formation, and glycan processing for glycoprotein maturation^8^.

Genome-scale metabolic models (GEMs) provide a comprehensive framework for the analysis of cellular metabolism at the systems level^9^. These constraint-based models represent the complete set of metabolic reactions within an organism and enable the prediction of metabolic fluxes under diverse conditions using flux balance analysis (FBA)^10^. The integration of viral biomass requirements into host metabolic models by constructing Viral Biomass Objective Functions (VBOFs) has become an effective strategy for investigating host-virus interactions and identifying potential antiviral targets^11^. This approach has been successfully applied to a range of viruses, including Chikungunya^11^, Dengue^11^, Zika^11^, SARS-CoV-2^12, 13^, and influenza^14^, thereby demonstrating its utility in identifying antiviral targets.

The VBOF approach quantifies the stoichiometric requirements for producing one complete virion. It includes all structural components, including nucleotides for the genome, amino acids for viral proteins (considering protein copy numbers), lipids for the envelope, and glycans for post-translational modifications. Integrating the VBOF as an objective function within the host metabolic model enables FBA to predict the metabolic state of infected cells. This method also facilitates identification of host reactions essential for viral production through systematic knockout analysis.

Previous studies have identified critical metabolic dependencies for viral replication, such as nucleotide metabolism, lipid biosynthesis, and amino acid transport^11, 15^. However, no comprehensive metabolic model currently exists for HMPV infection, which limits understanding of the specific host metabolic pathways required for HMPV replication and impedes identification of potential therapeutic targets. Recent pandemics, such as Coronavirus Disease 2019 (COVID-19), underscore the necessity of rapid response and real-time data monitoring. Furthermore, the virus is spreading rapidly, placing high-risk populations at increased vulnerability. Its potential adaptation to warmer climates poses a significant future public health risk^6, 16^.

In this study, a detailed VBOF for HMPV was constructed using experimental data from structural biology studies and related paramyxoviruses, supplemented by several assumptions. This VBOF was then integrated into *i*HBEC^1^, a human bronchial epithelial cell metabolic model that represents the primary target cell type for HMPV infection and was previously used for SARS-CoV-2 modeling^12, 13^. Systematic gene and reaction knockout analysis identified host metabolic genes and reactions essential for HMPV virion production. The results present a comprehensive map of host metabolic dependencies for HMPV replication and identify specific targets for antiviral drug development.

## 2 Results

### 2.1 HMPV Virion Composition and VBOF Construction

We constructed a comprehensive VBOF for HMPV that incorporates all major virion components.

#### 2.1.1 Genome stoichiometry

The nucleotide requirements for genome replication were determined directly from the genome sequence: 5,103 adenosine triphosphate residues (A), 3,402 uridine residues (U), 2,538 guanosine residues (G) and 2,307 cytidine residues (C).

#### 2.1.2 Protein stoichiometry

The copy number of the N protein was calculated based on the genome length of HMPV. HMPV has a genome length of 13,350 Kb and each N protein binds to 7 nucleotides, resulting in approximately 1,970 copies per virion. The 1,840 M protein copy number was estimated using a geometric surface area-based approach, as detailed in Section 5.1.2. A summary of the calculation is provided in Table 11.

The convex hull analysis showed that the YZ-plane projection of M Protein had the largest footprint area^17, 18^. This result supports the idea that the concave face acts as the binding surface of the membrane.

The copy numbers of the G, F, and SH proteins were computed using hexagonal packing analysis of surface glycoprotein spikes. These estimates varied depending on the spacing between spikes. The mean glycoprotein distribution on the virion surface was calculated as 416 total spikes, each with an average length of 11.5 nm and separated by an average gap of 8 nm. Based on these findings, the estimated numbers of glycoprotein subunits per virion are 693 F Protein, 139 G Protein, and 230 SH Proteins, resulting in a total of 1,062 glycoprotein subunits per virion.

The copy numbers of L and P proteins for HMPV were adopted from studies on Sendai virus, as both viruses belong to the same family. Accordingly, 50 L proteins and 300 P proteins were used for HMPV, mirroring the values reported for the Sendai virus^19, 20^.

The M2-1 protein in HMPV works as a tetramer, so 50 tetramers were assumed, matching the number of L proteins. This gave a total of 200 M2-1 protein subunits. The M2-2 protein was excluded from the initial calculations due to limited structural information. Later, its copy number was set to match both the highest (N protein) and lowest (N protein) protein copy numbers in separate tests. Changing the M2-2 copy number had no effect on viral growth, so the final Viral Biomass Objective Function (VBOF) was set to the lowest value of 50. In Figure 1, each protein copy number is assigned a confidence score according to the method and data source utilized. To improve the reliability of these confidence scores, future analyses should incorporate experimental data specific to HMPV as it becomes available.

**Figure 1.**
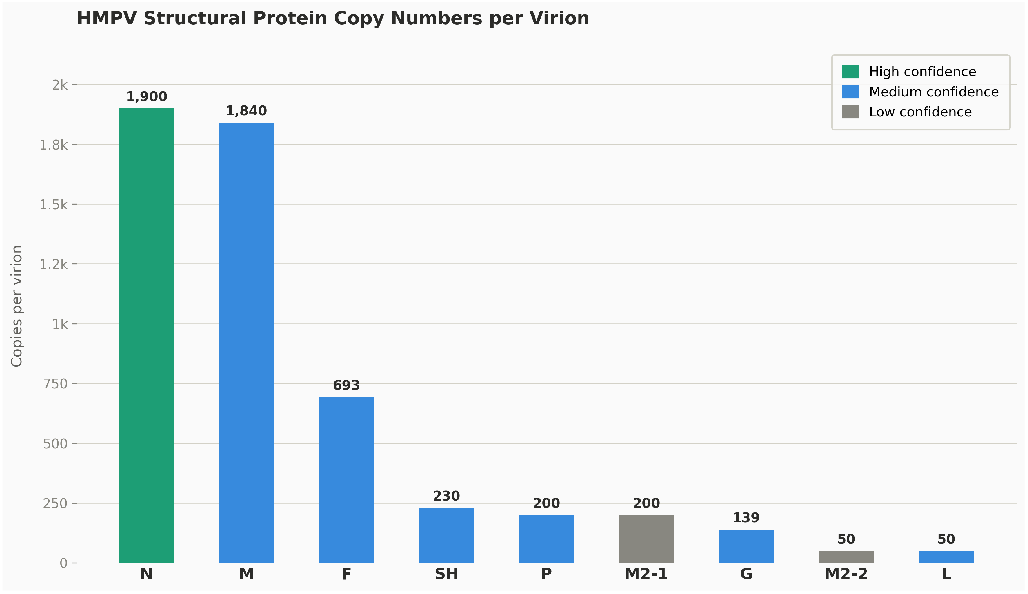
Protein copy numbers are classified into three confidence categories. High confidence indicates values calculated directly for HMPV using reliable methods. Medium confidence refers to values obtained from closely related viruses. Low confidence is assigned when values are assumed due to the absence of experimental data.

The total amino acid requirements per virion were calculated from protein copy numbers and amino acid sequence lengths, resulting in an estimate of 1,896,905 residues per virion, with leucine (189,025), serine (172,928), and alanine (153,879) being the most abundant.

#### 2.1.3 Lipid composition

The total number of lipid molecules in HMPV was estimated to be 388,587, based on its bilayer surface membrane area and the experimentally determined lipid composition of the Sendai virus, which was used as a proxy due to their shared classification within the Paramyxoviridae family^19^. Based on the lipid composition, the number of each individual lipid species was determined by multiplying the total number of lipids by the respective composition fraction Section 5.3. As shown in Table 1, the count of each lipid molecule is negative, indicating that these molecules originate from the host cell^21^.

**Table 1:**
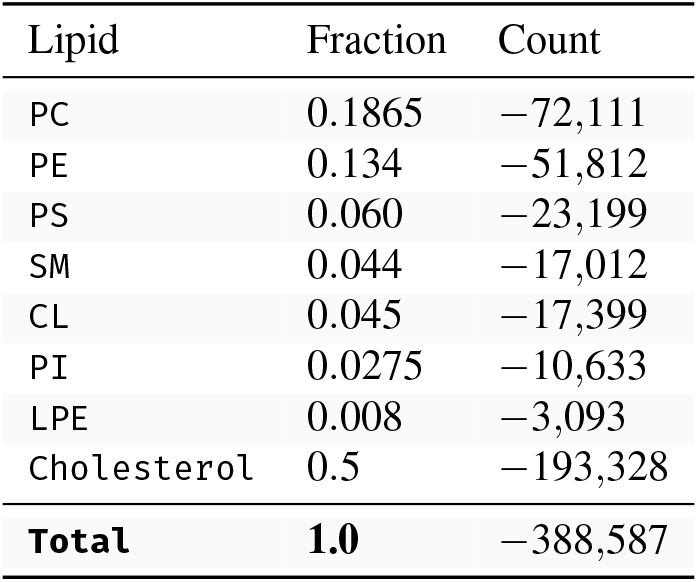
Lipid composition fractions and calculated number of molecules per virion.

#### 2.1.4 Glycan requirements

The F protein contains three established N-linked glycosylation sites that are consistently utilized during protein maturation. Given an estimated 695 F protein copies per virion, this results in 2,085 N-linked glycan structures present on the F protein.

The G protein contains between 3 to 6 N-linked glycosylation sites, which generate 444 distinct N-linked glycan structures. Additionally, O-linked glycosylation sites are present, with approximately 3,614 structures estimated to arise from 26 commonly utilized sites on each G protein. The comprehensive analysis revealed a total of 6,143 glycan structures per virion. To measure the metabolic needs for glycan biosynthesis *in silico*, we created models of typical structures for each glycan type. Then, we counted the individual monosaccharides for each glycan, based on its composition, and calculated the overall totals, shown in Table 13.

#### 2.1.5 Energy costs

The analysis also included the energy requirements for genomic RNA synthesis, tRNA charging, N- and O-linked glycan attachment, and ribosomal elongation, initiation, and termination. The total energy stoichiometries and their Biochemical, Genetical, and Genomical (BiGG) identifier (ID) are summarized in Table 2. These values were calculated based on VBOF metabolite counts, which include amino acids and glycans. The complete calculation methodology is provided in Section 5.4.3.

**Table 2:** Stoichiometry of energy metabolites in the VBOF of HMPV. Negative values indicate that a metabolite is consumed, and positive values mean it is produced.

| Metabolite | BiGG ID | Stoichiometry |
| --- | --- | --- |
| Adenosine triphosphate (ATP) | atp_c | −3,831,338 |
| Adenosine diphosphate (ADP) | adp_c | 3,831,338 |
| Adenosine monophosphate (AMP) | amp_c | 3,795,966 |
| Guanosine Triphosphate (GTP) | gtp_c | −3,795,984 |
| Guanosine diphosphate (GDP) | gdp_c | 3,795,984 |
| Pi | pi_c | 7,627,322 |
| H<sub>2</sub>O | h2o_c | −7,627,322 |
| H<sup>+</sup> | h_c | 7,627,322 |

#### 2.1.6 Complete VBOF

The model includes 44 metabolites, of which 38 are consumed and 6 are produced during virion assembly. Nucleotides, amino acids, glycans, and lipids constitute the primary metabolites in this VBOF. Energy cofactors, including ATP and GTP, represent the most heavily consumed resources. The top consumed metabolites available in the VBOF of HMPV are shown in Table 3. The breakdown products ADP, AMP, GDP, inorganic phosphate, and pyrophosphate are released as byproducts. The overall distribution of VBOF components by metabolite category is presented in Figure 2.

**Table 3:**
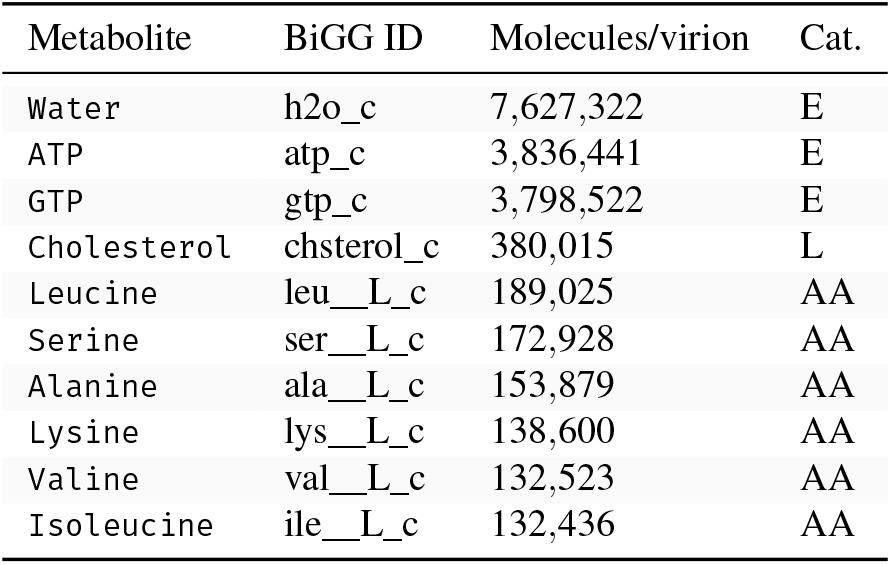
Top consumed metabolites in the VBOF of HMPV. Values represent molecules consumed per virion. Category (Cat.): E = Energy, AA = Amino acid, L = Lipid, G = Glycan.

**Figure 2.**
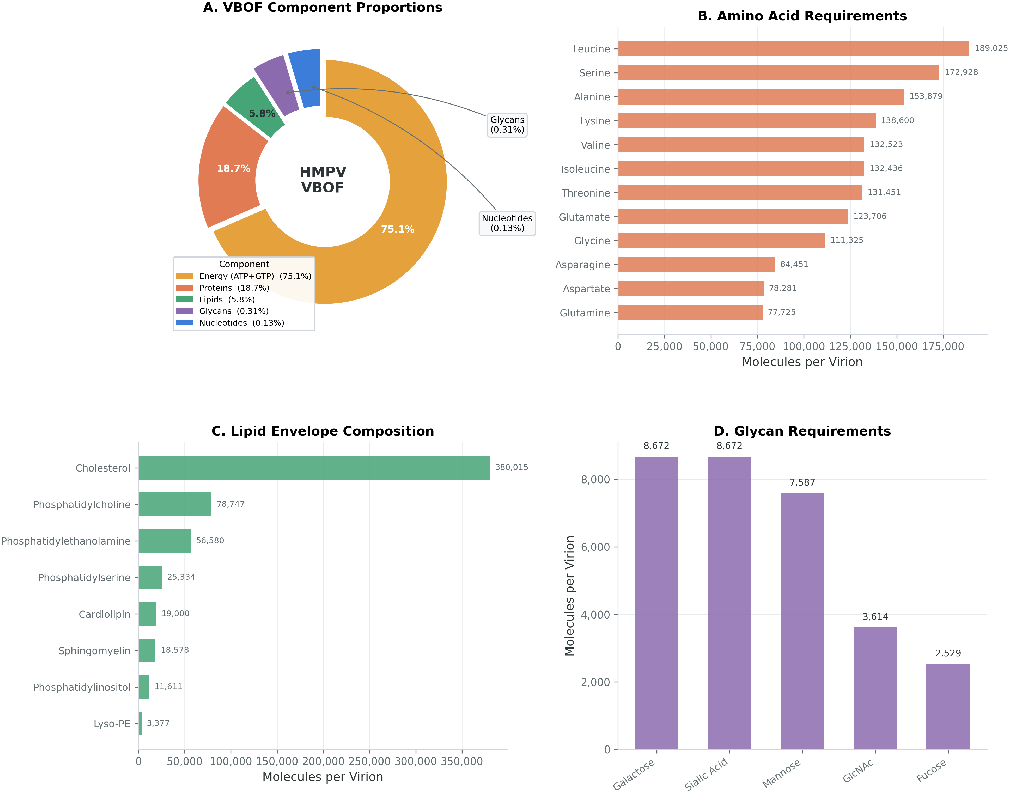
The composition of the VBOF in HMPV by metabolite category indicates that the primary energy demand, specifically ATP and GTP required for protein translation, constitutes the majority of VBOF flux. This is followed by contributions from amino acids, lipids, and glycan precursors. Nucleotides necessary for genome replication comprise a smaller yet essential fraction.

#### 2.1.7 Model Integration and Baseline Flux Analysis

The VBOF reaction was integrated into the human bronchial epithelial cell model *i*HBEC^1^. Of the 44 metabolites included in the VBOF reaction, 39 were mapped to the host model. The remaining five metabolites were not mapped, likely because the specialized lipid and glycosylation precursors they represent were absent from the reconstructed network. FBA using the HMPV_VBOF reaction as the objective yielded an optimal solution with an objective value of approximately 0.281 mmol*/*(g_DW_ · h). This result indicates that the host metabolic network can support HMPV virion production under the specified constraints.

The host biomass objective function (biomass_hbec) exhibited a flux rate of 0.234 mmol*/*(g_DW_ · h) under identical conditions. The substantially higher VBOF flux compared to the host biomass flux indicates the significant molecular complexity of the virion, especially its substantial energy and amino acid requirements. These results demonstrate that HMPV utilizes the metabolic capacity of epithelial cells to sustain virion production at a substantial rate.

### 2.2 Dual-Objective Gene Knockout Analysis

Host genes essential for HMPV production were identified by performing systematic single-gene knockouts, which were evaluated simultaneously against the VBOF and the host biomass objective in a dual-objective analysis. A total of 1,390 genes in the integrated model were systematically evaluated computationally.

#### Summary statistics

Gene knockouts were categorized according to their impact on host biomass flux (Table 4). In terms of viral production, 28 gene knockouts resulted in lethal flux loss (greater than 99 % reduction), one caused significant flux loss (50 % to 99 % reduction), and 1,361 had minimal effect (less than 10 % reduction). Regarding host viability, 34 gene knockouts were lethal to the host, and three resulted in a significant reduction of host flux.

**Table 4:** This summary presents the results of dual-objective single-gene knockout analysis, reporting the effects on both host biomass flux and viral VBOF flux for all 1,390 genes evaluated.

| Impact Category | Host Genes | Virus Genes |
| --- | --- | --- |
| Lethal (>99 % reduction) | 34 | 28 |
| Significant (50 % to 99 %) | 3 | 1 |
| Moderate (10 % to 50 %) | 0 | 0 |
| Minimal (<10 %) | 1,353 | 1,361 |
| **Total** | **1,390** | **1,390** |

#### Selective antiviral gene targets

Applying dual objective thresholds (viral production *<*50 %, host viability *>*80 %), we identified **two selective antiviral gene targets** (Table 5):

**Table 5:** Selective antiviral gene targets identified by dual-objective knockout analysis. Selectivity Score = Host Growth% −Virus Growth%.

| Gene | Function | Host | Virus | Score |
| --- | --- | --- | --- | --- |
| PGM3 | Phosphoacetylglucosamine mutase | 100 % | ≈0 % | 100 % |
| GNPNAT1 | GlcNAc-6-P acetyltransferase | 100 % | 6.13 % | 93.86 % |

1. **PGM_3_**(phosphoacetylglucosamine mutase): Knockout of this gene *in silico* resulted in complete loss of viral production (virus growth approximately 0 %) without detectable effects on host viability (100 % host growth retained). PGM_3_ catalyzes the conversion of N-acetylglucosamine-6-phosphate (GlcNAc-6-P) to GlcNAc-1-P, a key step in the hexosamine biosynthesis pathway that leads to UDP-GlcNAc synthesis.
2. **GNPNAT_1_**(glucosamine-6-phosphate N-acetyltransferase): Knockout of GNPNAT_1_ *in silico* reduced viral production to 6.13 % of baseline while preserving complete host viability (100 % host growth). GNPNAT_1_ catalyzes the acetylation of glucosamine-6-phosphate to generate GlcNAc-6-P, which is the step immediately upstream of PGM_3_ in the pathway.

#### Critical viral gene targets

Twenty nine genes were identified as critical viral targets, defined by viral production below 5 % irrespective of host effect. Among these, PGM_3_ and GNPNAT_1_ demonstrated the highest selectivity. Other critical targets include CMPK_1_ (UMP/CMP kinase; viral production 0 %, host growth 1.8 %) and a group of sterol biosynthesis genes (SPTLC_1_, SPTLC_2_, DHCR_24_, `MVD`, TM_7_SF_2_, CYP_51_A_1_, `NSDHL`, `MVK`, `EBP`, `LSS`, FDFT_1_, DHCR_7_, `SQLE`, SGMS_1_, `KDSR`, `HMGCR`, `FDPS`, SC_5_D, MSMO_1_), which are also lethal to the host. This finding highlights the mutual dependence on cholesterol and sphingolipids for both host cell membranes and the viral envelope.

### 2.3 Dual-Objective Reaction Knockout Analysis

All 1,769 reactions in the integrated model were systematically evaluated using single-reaction knockout analysis.

#### Summary statistics

A total of 57 reactions were lethal, and 4 were significant for viral production, while 67 reactions were lethal and 4 were significant for host viability (Table 6).

**Table 6:** Overview of dual-objective single-reaction knockout analysis involving 1,769 evaluated reactions.

| Impact Category | Host Rxns | Virus Rxns |
| --- | --- | --- |
| Lethal (>99 % reduction) | 67 | 57 |
| Significant (50 % to 99 %) | 4 | 4 |
| Moderate (10 % to 50 %) | 0 | 0 |
| Minimal (<10 %) | 1,698 | 1,708 |
| **Total** | **1,769** | **1,769** |

#### Selective antiviral reaction targets

Seven reactions satisfied the selectivity criteria, defined as viral production below 50 % and host viability above 80 %. Among these, one reaction was identified as a high-confidence target, exhibiting viral production below 10 % and host viability above 90 % (Table 7). The four highest ranking reactions are associated with the hexosamine biosynthesis and nucleotide sugar pathways:

**Table 7:**
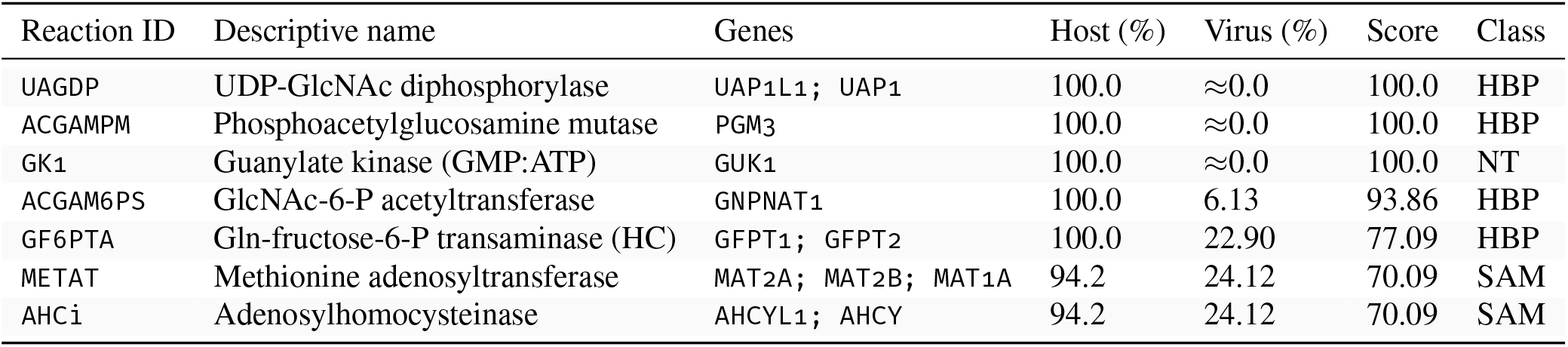
Selective antiviral reaction targets were identified using dual-objective knockout analysis and are presented in order of selectivity score. Reactions METAT and AHCi form part of the S-adenosylmethionine (SAM) cycle. Abbreviations: hexosamine biosynthesis pathway (HBP); nucleotide metabolism (NT) = nucleotide metabolism; S-adenosylmethionine (SAM) = S-adenosylmethionine cycle; High Confidence Target (HC) = High Confidence Target; Selective Target (S).

- **UDP-GlcNAc diphosphorylase (UAGDP)** (genes: `UAP1L1`, `UAP1`): Catalyzes the final step of Uridine Diphosphate (UDP)-GlcNAc synthesis (GlcNAc-1-P + UTP → UDP-GlcNAc + PPi). Knockout abolished vi-ral production while leaving host viability intact (100 % host growth, ≈0 % virus growth; selectivity score 100).
- **Phosphoacetylglucosamine mutase (ACGAMPM)** (gene: PGM_3_): Knockout abolished viral production with no host toxicity (100 % host, ≈0 % virus; score 100).
- **Guanylate kinase 1 (GK1)** (gene: `GUK1`): Knockout abolished viral production with no host toxicity (100 % host, ≈0 % virus; score 100). This reaction converts Guanosine Monophosphate (GMP) to GDP, which is essential for GTP production required for protein translation.
- **GlcNAc-6-P synthase (ACGAM6PS)** (gene: GNPNAT_1_): Knockout reduced viral production to 6.1 % of baseline with no host toxicity (score 94).
- **Glutamine-fructose-6-phosphate transaminase (GF6PTA)** (genes: `GFPT1`, `GFPT2`): The rate-limiting step of the hexosamine pathway. Knockout reduced viral production to 22.9 % of baseline with no host effect (score 77). This reaction is classified as a high-confidence target.
- **Methionine adenosyltransferase (METAT)** (genes: `MAT2A`, `MAT2B`, `MAT1A`): Knockout reduced viral production to 24.1 % of baseline with a minor host effect (host 94.2 %; score 70).
- **Adenosylhomocysteinase (AHCi)** (genes: `AHCYL1`, `AHCY`): Knockout reduced viral production to 24.1 % of baseline with a minor host effect (host 94.2 %; score 70).

#### Critical viral reaction targets

A total of 58 reactions resulted in greater than 95 % reduction in viral production (critical viral targets), comprising the seven previously identified selective targets and 51 additional reactions primarily associated with cholesterol and sphingolipid biosynthesis, which are also essential for host viability.

### 2.4 Visualization of Dual-Objective Analysis

The dual-objective scatter plot (Figure 3) illustrates the relationship between host and viral growth upon each gene knockout. Ideal drug targets are located in the upper-left quadrant, which corresponds to high host survival and low viral survival. The close clustering of most knockouts along the diagonal representing preserved host viability and minimal viral growth, with the notable upper-left outliers PGM_3_ and GNPNAT_1_, underscores the specificity of the hexosamine pathway in viral glycan biosynthesis.

**Figure 3.**
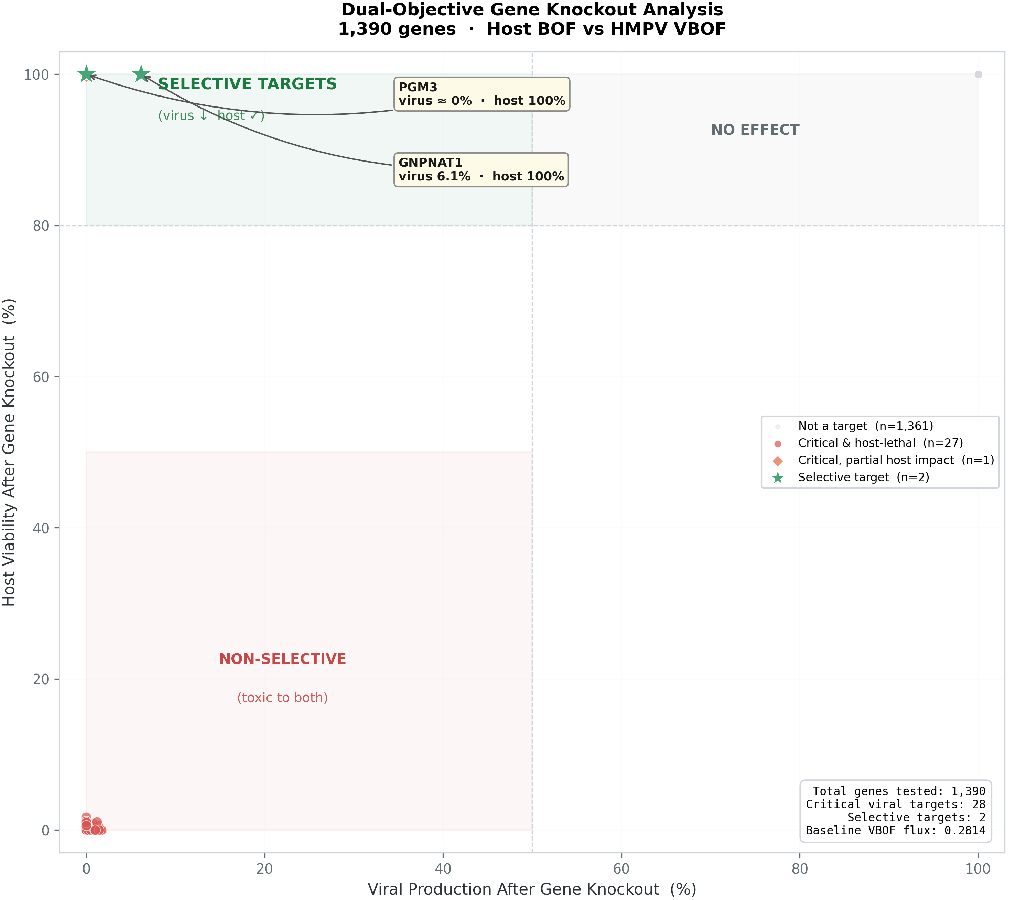
The dual-objective scatter plot shows gene-knockout analysis, where each point corresponds to a single gene knockout and plots host growth (%) against viral production (%). The optimal drug target region, characterized by high host survival and minimal viral survival, is indicated in the upper-left corner. PGM_3_ and GNPNAT_1_ are identified as the most selective antiviral targets.

Figure 4 presents a detailed comparison of the most selective targets, highlighting differences in viral and host growth suppression.

**Figure 4.**
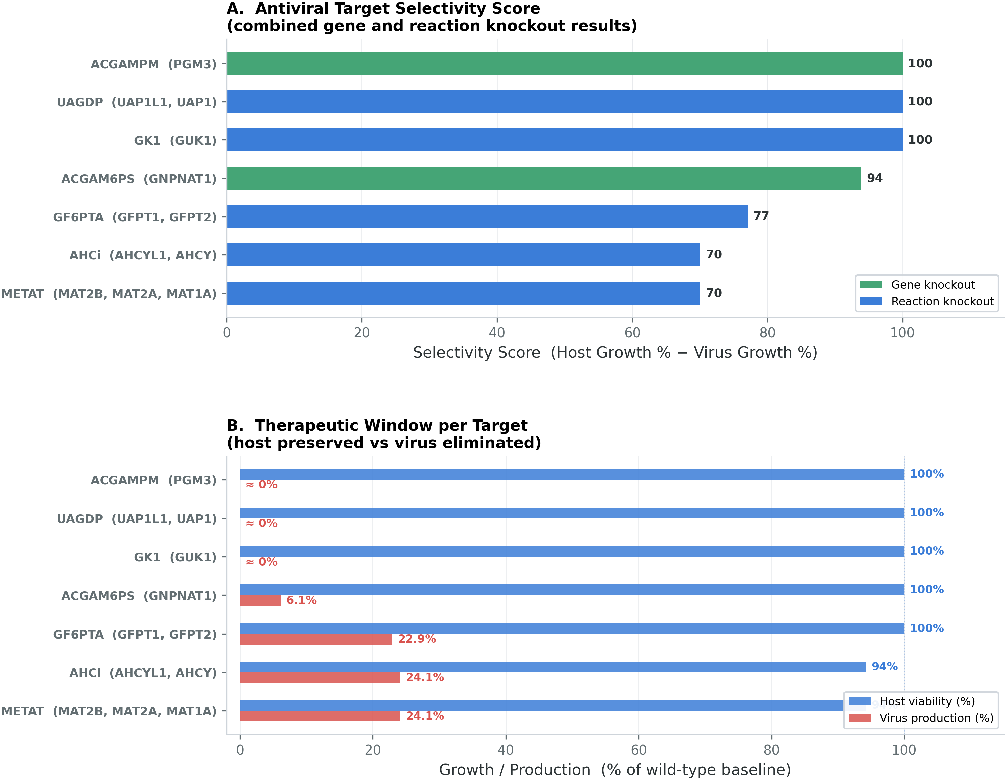
Top selective antiviral gene and reaction targets ranked by selectivity score (Host Growth % − Virus Growth %). All top-ranked targets belong to the hexosamine biosynthesis pathway (PGM_3_, GNPNAT_1_, UAP_1_/UAP_1_L_1_, GFPT_1_/GFPT_2_) or nucleotide metabolism `(GUK1)`.

## 3 Discussion

### 3.1 The HBP as the Primary HMPV Metabolic Vulnerability

Previous experimental studies have established that the hexosamine biosynthesis pathway is functionally important for HMPV replication^8^. Consistent with this, the primary finding of the present study identifies the **hexosamine biosynthesis pathway (HBP)** as the dominant selective metabolic vulnerability for HMPV replication. Four of the seven selective antiviral reaction targets (UAGDP, ACGAMPM, ACGAM6PS, GF6PTA) and both selective gene targets (PGM_3_, GNPNAT_1_) belong to this pathway (Figure 4).

The HBP converts fructose-6-phosphate into UDP-GlcNAc through four enzymatic steps: (1) GFPT_1_GFPT_2_(GF6PTA) converts fructose-6-P + glutamine to glucosamine-6-P; GNPNAT_1_ (ACGAM6PS) acetylates glucosamine-6-P to GlcNAc-6-P; (3) PGM_3_(ACGAMPM) isomerizes GlcNAc-6-P to GlcNAc-1-P; and (4) UAP_1_UAP_1_L_1_(UAGDP) condenses GlcNAc-1-P with UTP to form UDP-GlcNAc, the terminal product.^22^ All four steps are selectively lethal for HMPV production without host toxicity, demonstrating that the entire HBP is a valid antiviral target pathway.

The biological basis for this dependence is well established: UDP-GlcNAc serves as the essential sugar-nucleotide donor for O-GlcNAc modifications that densely coat the HMPV G protein^23^. Each HMPV virion contains approximately 45 O-linked glycosylation sites per G protein monomer and 250 G protein copies, resulting in a requirement of over 7,300 UDP-GlcNAc molecules for G protein O-glycosylation alone^24^. These mucin-like O-glycans on the G protein are functionally important for cell attachment and immune evasion^8^. In contrast, host cells maintain UDP-GlcNAc primarily for protein O-GlcNAcylation of transcription factors and structural proteins^25^. This requirement can be satisfied at a lower HBP flux than is necessary for viral glycan production^26^.

The enzymes PGM_3_and GNPNAT_1_represent promising drug candidates. Knockout of either gene resulted in complete elimination of viral production while maintaining 100 % host cell viability, with selectivity scores of 100 and 97.4, respectively. Germline mutations in PGM_3_in humans cause a rare primary immunodeficiency characterized by hyper-IgE syndrome^27^. This finding suggests that partial inhibition, rather than complete loss, may be more clinically tolerable. The upstream enzyme GFPT_1_GFPT_2_(GF6PTA) is also considered a high-confidence target (virus 9.6 %, host 100 %). The glutamine analog 6-Diazo-5-oxo-L-norleucine (DON) is already known to inhibit Glutamine—fructose-6-phosphate aminotransferase (GFPT)^28^, which provides a pharmacological entry point.

### 3.2 Guanylate Kinase (GUK1/GK1) as a Conserved Antiviral Target

The guanylate kinase reaction (GK1, catalyzed by `GUK1`) was identified as the third high-selectivity reaction target, exhibiting a selectivity score of 100 (viral growth approximately 0 %, host growth 100 %). `GUK1` catalyzes the phosphorylation of GMP to GDP^29^, thereby supplying the GTP pool necessary for RNA polymerase elongation and ribosomal translocation during viral protein synthesis.

Previous studies identified `GUK1` as a robust antiviral target against SARS-CoV-2 across multiple viral variants^12, 13^. These findings indicate that guanylate kinase constitutes a pan-viral metabolic vulnerability in RNA viruses with large genomes that require substantial GTP supply. The conservation of this finding across diverse RNA viruses, including SARS-CoV-2 and HMPV, suggests that `GUK1` inhibitors may serve as the foundation for broad-spectrum antiviral strategies.

### 3.3 S-Adenosylmethionine (SAM) Cycle Targets

METAT and AHCi, both components of the SAM cycle^30^, represent selective antiviral targets that exhibit modest toxicity to host cells (host growth 94.2 %, virus growth 11.1 %, score 83.1). SAM serves as the universal methyl donor for a wide range of cellular methylation reactions^31^, including the 5’ capping of messenger ribonucleic acid (mRNA)^32^. HMPV, like other negative-sense RNA viruses, encodes its own cap-methyltransferase activity within the L protein; however, this enzymatic activity remains dependent on the host SAM pool^33^.

The predicted partial impact on the host indicates that complete *in silico* inhibition of the SAM cycle may not be well tolerated. However, combination strategies targeting both SAM synthesis and HBP enzymes could yield additive antiviral effects at sub-toxic doses of each agent.

### 3.4 Sterol and Sphingolipid Biosynthesis: Shared Host-Virus Dependencies

The largest group of critical viral targets, comprising 21 genes, is associated with the cholesterol and sphingomyelin biosynthesis pathways (`HMGCR`, `MVK`, FDFT_1_, `LSS`, CYP_51_A_1_, `NSDHL`,DHCR_7_, DHCR_24_, `SQLE`, SGMS_1_, `KDSR`, among others). Although these pathways are essential for HMPV envelope formation, with sphingomyelin (15 %) and cholesterol (10 %) together comprising 25 % of the viral envelope, they are also crucial for maintaining host cell membrane integrity. As a result, these targets are considered critical for viral replication but lack selectivity, which limits their potential as therapeutic interventions. However, strategies involving partial inhibition or the use of membrane-targeting antivirals, such as lipid raft disruptors, may merit further investigation. Statins, which inhibit `HMGCR`, have demonstrated antiviral activity in cell culture models of related paramyxoviruses. However, clinical evidence remains inconclusive.

### 3.5 Comparison with Metabolic Models of Other Respiratory Viruses

A recurring finding in GEM-based antiviral target studies is the central importance of nucleotide metabolism, with a specific emphasis on the purine pathway^1, 12, 13, 14^. The identification of `GUK1` as a selective HMPV target parallels findings in SARS-CoV-2, reinforcing the concept that purine metabolism represents a broadly exploitable antiviral vulnerability in enveloped RNA viruses.

### 3.6 Limitations and Future Directions

Several limitations must be acknowledged when interpreting these results. First, the model provides only a static representation of cellular metabolism under standard growth conditions. *In vivo*, HMPV infection induces dynamic alterations in host cell metabolism, such as activation of interferon signaling, increased redox stress, and reorganization of central carbon metabolic pathways. Future studies should integrate time-resolved transcriptomic constraints to develop condition-specific models of infected cells^1, 14^.

Second, the single-gene and single-reaction knockout framework fails to account for genetic redundancy or compensatory metabolic mechanisms that may be activated in infected cells. Combinatorial knockout and flux variability analyses would yield a more comprehensive assessment of metabolic network robustness.

Third, protein copy numbers and other structural numbers are not experimentally based because HMPV-specific structural data are currently unavailable. More accurate values from future cryo–electron microscopy (cryoEM) studies would enhance model fidelity.

Fourth, the model omits dynamic viral-host signaling and immune responses. Host-directed therapies identified in this study should be evaluated within the context of the host immune response, as these pathways may also be influenced by the same metabolic processes.

Future research priorities are as follows: (1) experimental validation of PGM_3_, GNPNAT_1_, UAP_1_, and `GUK1` as antiviral targets using Small interfering RNA (siRNA) knockdown and selective small-molecule inhibitors in HMPV-infected bronchial epithelial cells; (2) integration of transcriptomic data from HMPV-infected cells to construct condition-specific metabolic models; (3) systematic screening of existing HBP inhibitors, including GFPT inhibitors such as DON, against HMPV in cell culture; (4) extension of dual-objective analyses to combination therapies; and (5) application of this framework to RSV, a closely related paramyxovirus, to identify both shared and virus-specific targets.

## 4 Conclusion

We introduce the first genome-scale metabolic model of HMPV–host interaction, developed by integrating a comprehensive VBOF for HMPV with the human bronchial epithelial cell model *i*HBEC^1^. The VBOF defines the full stoichiometric requirements for HMPV virion production across five metabolite categories: nucleotides (genome replication), amino acids (nine viral proteins, 1,682,580 residues per virion), energy (translation and replication), lipids (envelope), and glycans (9,162 glycosylation events per virion).

Dual-objective knockout analysis of 1,390 genes and 1,769 reactions in the integrated model identified two selective antiviral gene targets (PGM_3_ and GNPNAT_1_) and seven selective reaction targets, primarily within the hexosamine biosynthesis pathway (HBP). Knockout of both PGM_3_ and GNPNAT_1_ completely abolished HMPV production while maintaining 100 % host cell viability. This selectivity profile is highly advantageous for antiviral drug development. This selectivity arises from the exceptionally high demand for UDP-GlcNAc generated by extensive O-glycosylation of the HMPV G protein, which surpasses the host cell’s own UDP-GlcNAc requirements.

Guanylate kinase (`GUK1`) was identified as a third high-selectivity target, consistent with its identification as a robust target in SARS-CoV-2 models^12, 13^. This finding supports the existence of conserved vulnerabilities in nucleotide metabolism among RNA viruses.

Given the absence of approved vaccines or antiviral therapies for HMPV, the host metabolic targets identified in this study, particularly PGM_3_, GNPNAT_1_, and GUK_1_, represent promising candidates for further experimental and therapeutic investigation. The computational framework and integrated metabolic model developed in this study are applicable to other respiratory viruses and are freely accessible to facilitate future research. Figure 5 illustrates the workflow developed for creating a VBOF, integrating it into the host model, and conducting subsequent analyses.

**Figure 5.**
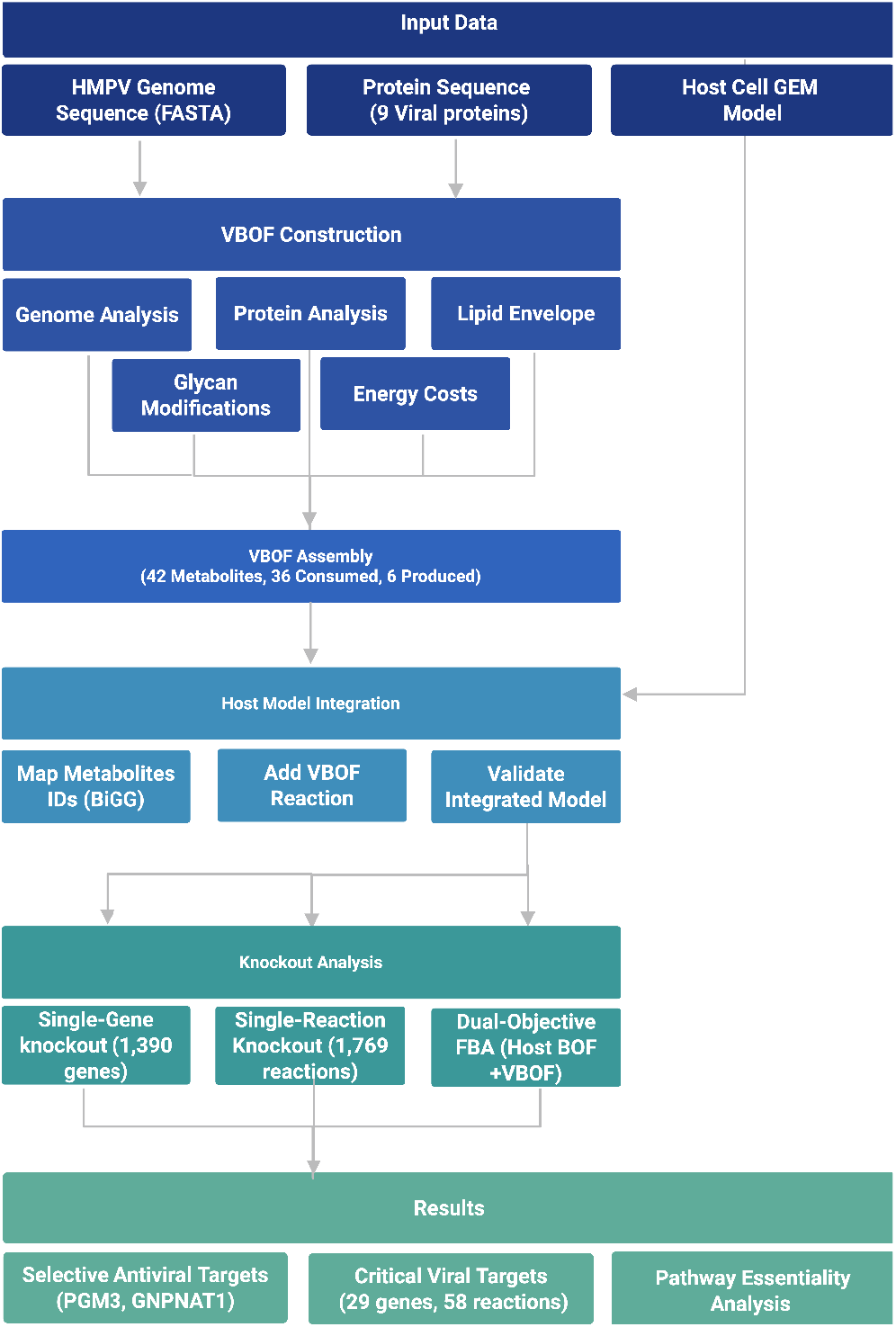
The computational pipeline for construction of VBOF in HMPV and antiviral target identification. The workflow includes genome and protein sequence retrieval, VBOF assembly, host model integration, and dual-objective FBA-based knockout analysis. Figure created with BioRender (BioRender.com).

## 5 Materials and Methods

### 5.1 Protein Copy Number Determination

Protein copy numbers per virion (Table 8) were determined using a combination of direct calculations, experimental estimates from the closely related RSV, and informed assumptions for proteins lacking experimental data. Each estimate was assigned a confidence level (high, medium, or low) as shown in Figure 1.

**Table 8:** HMPV viral protein composition and copy numbers per virion. Protein lengths and copy numbers were derived from experimental structural studies (see Section 5).

| Protein | Function | Length (aa) | Copies /virion | aa/virion |
| --- | --- | --- | --- | --- |
| N | Nucleoprotein | 394 | 1,900 | 751,358 |
| M | Matrix protein | 254 | 1,840 | 467,360 |
| F | Fusion protein | 539 | 693 | 373,527 |
| P | Phosphoprotein | 294 | 300 | 88,200 |
| G | Attachment glycoprotein | 236 | 139 | 32,804 |
| M2-1 | Transcription processivity | 187 | 200 | 37,400 |
| L | Large polymerase protein | 2,005 | 50 | 100,250 |
| SH | Small hydrophobic protein | 183 | 232 | 42,456 |
| M2-2 | Regulatory accessory protein | 71 | 50 | 3,550 |
| **Total** |  |  | 5,676 | 1,896,905 |

#### 5.1.1 Estimation of N Protein copy number

The N protein encapsidates the viral RNA genome, with each N protein protomer binding a specific number of nucleotides. The copy number of the N protein can be directly inferred from the genome length and the binding stoichiometry of the N-RNA interaction:

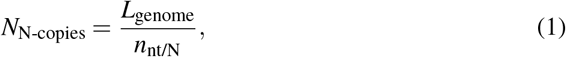

where *L*_genome_ is the total number of nucleotides in the viral genome and *n*_nt/N_ is the number of nucleotides bound per N protomer.

The HMPV genome consists of a single-stranded, negative-sense RNA of approximately 13,350 nucleotides. The number of nucleotides bound per N protein protomer was determined from the crystal structure of the closely related RSV nucleoprotein-RNA complex. In this structure, RNA wraps around the protein ring, with each N protein subunit contacting seven nucleotides^34^.

This value of *n*_nt/N_ = 7 nucleotides per N protomer was confirmed by subsequent structural studies^35^. The application of RSV N protein binding stoichiometry to HMPV is supported by the substantial structural conservation reported between RSV and HMPV nucleoproteins^36^.

For virions containing a single genome copy, the N protein copy number is approximately 1,900. This value increases in virions that incorporate multiple genome copies. For instance, filamentous RSV particles typically contain two genome copies, whereas spherical particles range from 1 to 9 genome copies^37^.

#### 5.1.2 Estimation of M Protein copy number

The M protein of HMPV lines the inner leaflet of the viral lipid bilayer and functions as the primary structural scaffold connecting the viral envelope to the Ribonucleoprotein (RNP) complex^38^. To date, no experimental measurement of the M protein copy number in HMPV virions has been reported. This study estimates copy number of M protein using a geometric surface-area-based approach, using HMPV specific structural data when available and incorporating parameters from Human respiratory syncytial virus (hRSV) when HMPV specific data are unavailable. The copy number of a matrix protein that covers the inner membrane surface of a viral particle can be estimated using the following geometric relationship:

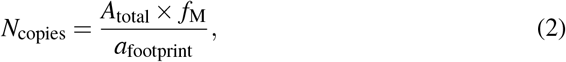

where *A*_total_ is the total membrane surface area of the virion, *f*_M_ is the fraction of the membrane covered by the M protein layer, and *a*_footprint_ is the area occupied by one M protein unit (dimer or monomer) on the membrane surface.

This approach was first applied to viral protein stoichiometry of Human immunodeficiency viruses (HIV) Gag protein copy numbers by calculating the number of hexameric unit cells that fit on the spherical inner surface of the virion^39^. It should be noted that geometric estimates assume uniform, gap-free packing and therefore not represent 100 % accurate results.

The mean diameter of spherical HMPV particles was taken as 209 nm (radius *r* =104.5 nm). Based on this diameter, the total membrane surface area for a spherical particle is:

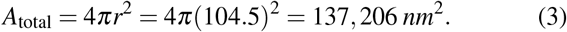

To date, no cryo-electron tomography studies have measured M coverage in intact HMPV particles. Therefore, M coverage values were adapted from cryo-ET measurements of hRSV. The surface area of the membrane covered with M was calculated to be *∼*24 % for spherical hRSV virus^37^. For the spherical HMPV particle, *f*_M_ = 0.24, giving a M-covered area of:

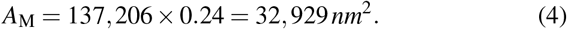

The use of hRSV coverage values for HMPV is justified by several considerations: (i) HMPV and hRSV are classified together in the *Pneumovirinae* subfamily and share the same three morphological classes^40^, and (ii) their M protein share 38 % amino acid sequence identity^38^.

##### M Protein dimer footprint determination

The footprint of the HMPV M protein dimer on the membrane surface was determined from the crystal structure coordinates (Protein Data Bank (PDB) 4LP7)^41^ using the following procedure:

a. The biological assembly of PDB 4LP7, comprising chains A and C, was obtained from the Research Collaboratory for Structural Bioinformatics (RCSB) Protein Data Bank^41^.
b. The asymmetric unit consists of four chains (A, B, C, D).
c. Figure 6 shows chains A and C, which constitute the biologically relevant dimer^38^.
d. Atomic coordinates were extracted using BioPython^42^, and the analysis included all non-hydrogen atoms.

**Figure 6.**
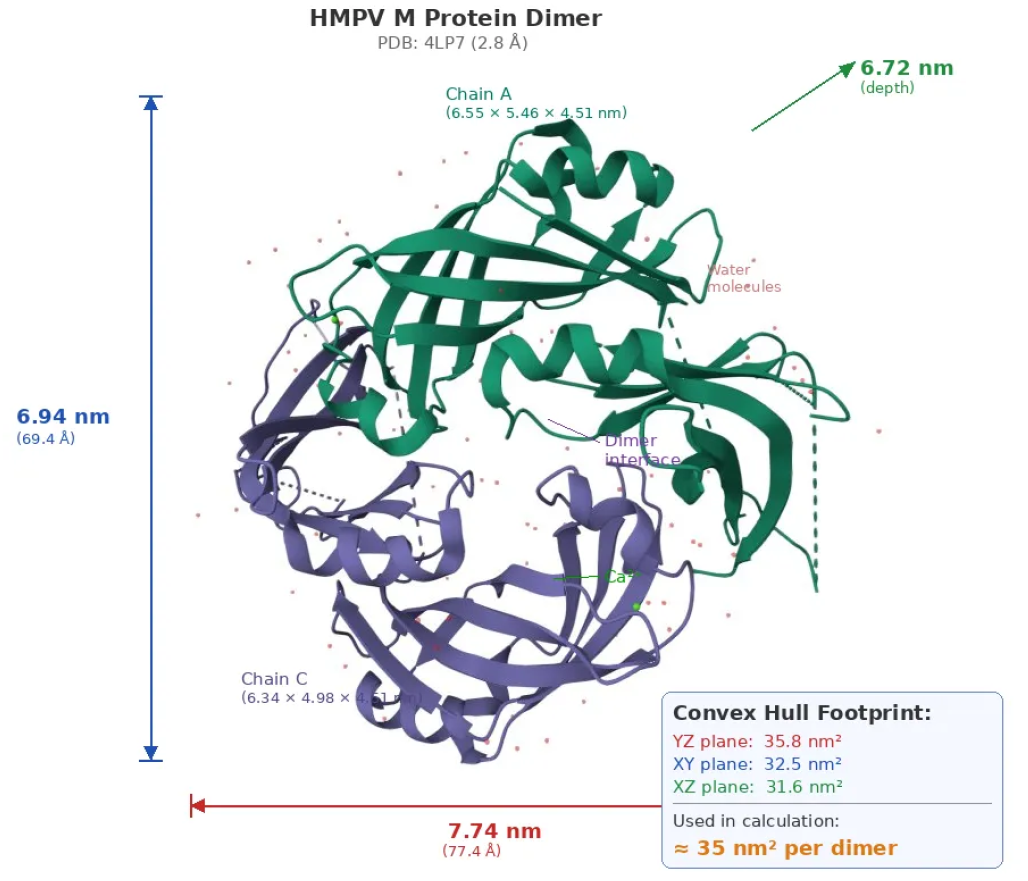
Structural dimensions of the HMPV M protein dimer^41^ resolution.

##### Bounding box dimensions

The spatial extent of the dimer was calculated based on the range of Cartesian coordinates along each axis as shown in Table 9. These values correspond to the maximum bounding box dimensions and served as an initial estimate.

**Table 9:**
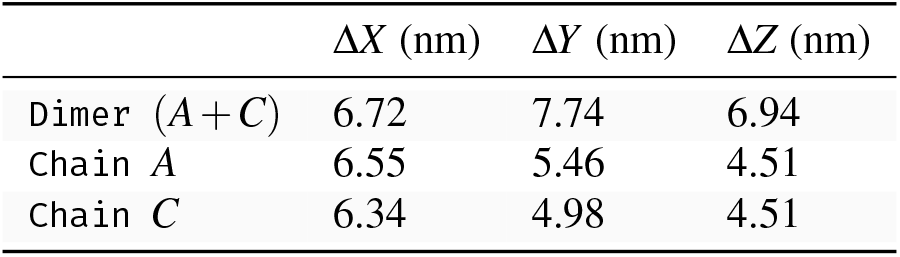
Bounding box dimensions of the HMPV M dimer biological assembly (PDB 4LP7, chains *A*+*C*). Individual chain dimensions are provided for comparison.

##### Convex hull projection

A more accurate footprint that reflects the protein’s irregular shape, without assuming a packing fraction, was obtained by projecting the atomic coordinates onto each of the three cardinal planes (*XY, XZ, YZ*). The convex hull, defined as the minimum enclosing polygon, of the resulting two-dimensional point set was then computed using SciPy’s `ConvexHull`^43^ implementation (see Table 10).

**Table 10:** Convex hull areas of the HMPV M dimer projected onto three cardinal planes.

| Projection plane | Area (nm<sup>2</sup>) | Interpretation |
| --- | --- | --- |
| YZ-plane | 35.8 | Membrane-facing face |
| XY-plane | 32.5 | Lateral view |
| XZ-plane | 31.6 | End view |

**Table 11:** Geometric estimation of the HMPV M protein copy number for a spherical virion with a mean diameter of 209 nm.

| Step | Operation | Value | Source |
| --- | --- | --- | --- |
| Radius | $209/2$ | 104.5 nm | 7 |
| Surface area | $4\pi r^2$ | 137,206 nm<sup>2</sup> | — |
| M-covered area | $\times 0.24$ | 32,929 nm<sup>2</sup> | 37 |
| Dimer count | $\div 35.8 \text{ nm}^2$ | 920 dimers | 38 |
| **Monomer count** | $\times 2$ | **1,840** | — |

##### Selection of the membrane-facing projection

There are two possible orientation of the M protein dimer, the concave (positively charged) face toward the lipid and the convex face toward the lipid. The concave-face-down orientation produced a substantially higher fitting correlation (*C* = 0.94) compared to the alternative (*C* = 0.88), thereby demonstrating that the concave face serves as the membrane-binding surface^38^.

The concave face corresponds to the largest projected area of the dimer, the *YZ*-plane projection at 35.8 nm^2^, consistent with the bowl-shaped geometry, which presents a larger cross section when viewed from the membrane normal (Table 10). This value was adopted as the dimer footprint in Equation (2). Application of Equation (2) with *r* = 104.5 nm, *f* = 0.24 nm^2^, and *a* = 35.8 nm^2^ yielded:

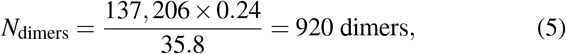

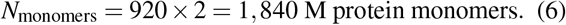

The calculation is summarized in Table 11.

#### 5.1.3 Estimation of Glycoprotein (F, G and SH) copy number

HMPV, like RSV, possesses three principal surface glycoproteins: the fusion protein (F), attachment glycoprotein (G), and small hydrophobic protein (SH)^44, 45^. While experimental studies have not yet established the precise copy numbers or stoichiometric ratios of these proteins on HMPV virions, electron microscopy analyses of RSV demonstrate densely packed glycoprotein spikes distributed across the viral envelope surface^46^. Due to the close phylogenetic relationship between HMPV and RSV within the Pneumoviridae family, these structural parameters serve as a proxy for estimating glycoprotein abundance on HMPV particles.

##### Glycoprotein Spike Dimensions

Electron microscopy studies of RSV demonstrate that individual glycoprotein spikes possess a width of approximately 11 nm to 12 nm (average: 11.5 nm) and are separated by gaps ranging from 6 nm to 10 nm (average: 8 nm), resulting in a thistle-like appearance of the virion surface ^47^. These measurements were subsequently used as a proxy for HMPV spike geometry.

The center-to-center distance between adjacent spikes was calculated as follows:

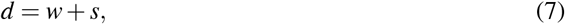

where *w* is the spike width and *s* is the spacing between spikes. Therefore, we have

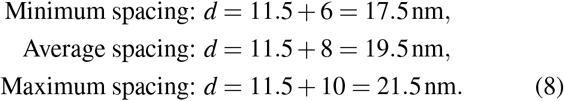

##### Justification for Hexagonal Packing Model

Hexagonal close packing was used as the basis for estimating glycoprotein density on the virion. This assumption is supported by three independent lines of evidence: (i) cryogenic electron tomography (cryo-ET) of RSV has directly revealed honeycomb-like hexagonal arrays of glycoprotein spikes on pleomorphic virions, with a mean inter-spike spacing of 74 angstrom(AA) at the hexagonal vertices^46^., (ii) hexagonal lattice formation by surface glycoproteins has been demonstrated on the related influenza C virus through subtomogram averaging^48^, establishing hexagonal packing as a general feature of enveloped virus glycoprotein, and (iii) the hexagonal arrangement represents the mathematically optimal configuration for packing identical objects on a two-dimensional surface, thereby achieving maximal packing density of 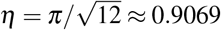, as proven by Lagrange’s Four-Square Theorem (1773) for lattice packings^49^ and by Fejes Tóth for all arrangements^50^. This optimality result supports selecting hexagonal geometry as the preferred packing model.

On a spherical surface, a perfect hexagonal lattice necessarily incorporates exactly 12 pentagonal defects. This requirement is explained by Caspar–Klug Quasi-Equivalence Theory^51, 52, 53^. For the spike counts estimated in this study (343 to 517), these 12 defects represent approximately 2 % to 3 % of lattice sites. However, since pentagonal sites still accommodate a spike, differing only in local coordination number, the effect on the overall density estimate is substantially lower than this fraction and is negligible compared to the uncertainty introduced by the measured variation in inter-spike spacings.

##### Spike Density Calculation

In a hexagonal packing with nearest neighbor spacing *d*, each spike occupies a Voronoi cell, which is a regular hexagon with area given by:

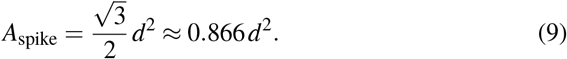

Subsequently, the total number of spike positions on the virion surface was calculated as follows:

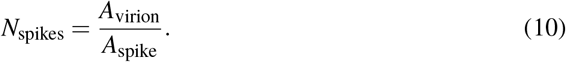

Using the calculated virion surface area from Equation (3) and different spacings from Equation (8), we have as follows:

- **Minimum spacing (**17.5 **nm)**

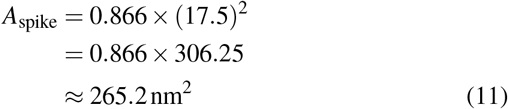

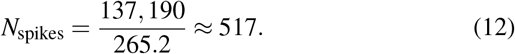
- **Average spacing (**19.5 **nm)**

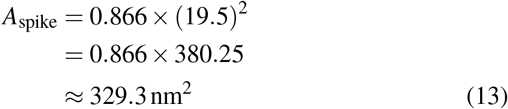

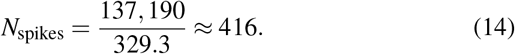
- **Maximum spacing (**21.5 **nm)**

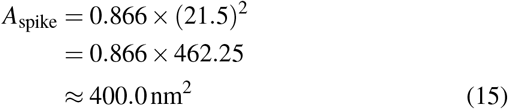

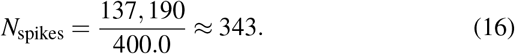

##### Glycoprotein Ratio Determination

To date, there have been no experimental reports quantifying the F:G:SH stoichiometry on HMPV or RSV virions. Consequently, we estimated the relative abundance of each glycoprotein using three independent lines of evidence:

i. **Transcription gradient**- In paramyxoviruses, the viral RNA-dependent RNA polymerase (RdRp) (which is an enzyme used by many RNA viruses to copy their RNA genomes) initiates transcription at a single 3^*′*^ promoter and sequentially transcribes each gene, with a probability of polymerase disengagement at each gene junction. This process generates a gradient of mRNA abundance that decreases from the 3^*′*^ to the 5^*′*^ end of the genome^54^. In the HMPV genome (3^*′*^-N-P-M-F-M2-SH-G-L-5^*′*^), the F gene is located upstream of both SH and G^55^, which predicts higher F mRNA and, consequently, greater F protein abundance relative to proteins SH and G.
ii. **Functional essentiality**- The F protein is the sole HMPV glycoprotein required for both cellular attachment via RGD-binding integrins (which are a subgroup of integrin receptors that recognize the short amino acid sequence) and membrane fusion, and it is the primary inducer of neutralizing antibodies^56, 57^. In contrast, G-deleted recombinant HMPV remains viable *in vitro* and can replicate in the upper respiratory tract of non-human primates, suggesting that the G protein serves as an auxiliary attachment factor^55, 56^. Similarly, SH-deleted recombinant HMPV, a genetically engineered HMPV virus lacking the SH gene, replicates with slightly reduced efficiency in both hamster and non-human primate models, indicating that the SH protein is not essential for viral replication^55, 58^.
iii. **Relative protein abundance**- F and G proteins have been identified as the main protein constituents of the HMPV envelope^56^, while the SH protein is present at comparatively low level and is considered functionally dispensable. These findings support the conclusion that F protein is the most abundant surface glycoprotein, followed by G protein, with SH protein representing a minor component.

Based on these findings, a stoichiometric ratio was adopted as follows:

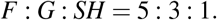

Therefore, total ratio sum is

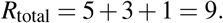

##### Individual Glycoprotein Copy Number Calculation

The copy number of each glycoprotein was calculated using:

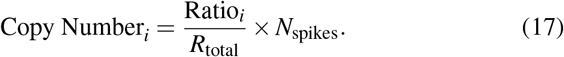

Accordingly, the glycoprotein spike number per HMPV was estimated using different spike calculations based on spacings defined in Equation (12), Equation (14), and Equation (16), with an F:G:SH ratio of 5:3:1. The results are summarized in Table 12.

**Table 12:**
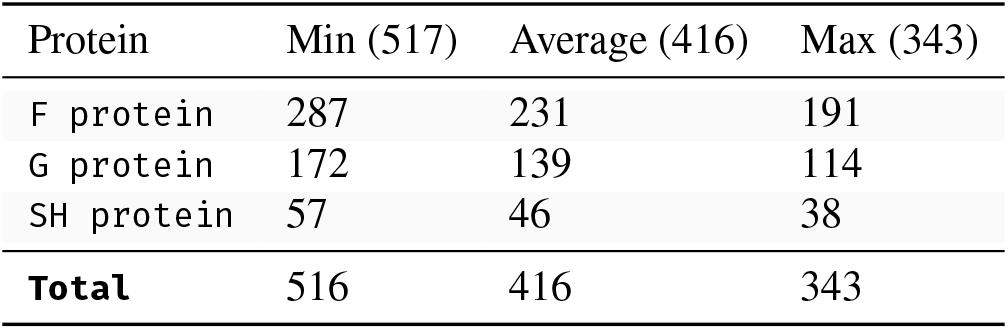
The estimated number of glycoprotein spikes per HMPV virion based on the F:G:SH ratio of 5:3:1. Rounding in the Min column (517) resulted in a summation discrepancy, yielding a total of 516.

**Table 13:** The total monosaccharide requirements are determined based on N-linked and O-linked glycan structures present in the HMPV virion.

| Monosaccharide | Total Count |
| --- | --- |
| GlcNAc | 10,116 |
| Man | 7,587 |
| Gal | 8,672 |
| Neu5Ac | 8,672 |
| Fuc | 2,529 |
| GalNAc | 3,614 |

##### Glycoprotein Subunit Numbers

Given the oligomeric structure of HMPV glycoproteins, the number of individual protein subunits per virion was determined based on spike counts.

The F protein forms a homotrimer composed of disulfide-linked F1–F2 heterodimers, which constitutes the functional unit responsible for mediating membrane fusion^44, 56^. The G protein of HMPV predominantly exists as a monomer in its soluble form and functions as an intrinsically disordered, heavily glycosylated polymer^45, 59^. The precise oligomeric state of HMPV SH protein remains undetermined. However, based on analogy with the closely related RSV SH protein, which forms pentameric ion channels as demonstrated by solution NMR and analytical ultracentrifugation^60^, HMPV SH protein was assumed to assemble as pentamers for this calculation. Additionally, HMPV SH protein has been shown to form higher-order oligomers consistent with viroporin function^58^. For the average spacing case, the protein subunits are as follows:

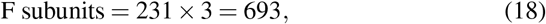

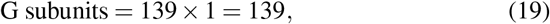

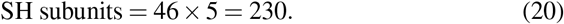

Therefore, the estimated total number of glycoprotein subunits per virion is 693 + 139 + 230 = 1,062.

### 5.2 Estimating Glycan Requirements for the HMPV Virion

The carbohydrate requirements for the VBOF of HMPV were estimated by modeling the glycosylation of the two major envelope glycoproteins of HMPV: the fusion (F) protein and the attachment (G) protein. The number of glycan structures associated with each protein was calculated, and the total monosaccharide composition of the virion was subsequently derived from representative glycan structures.

#### 5.2.1 F Protein Glycosylation

The HMPV F protein possesses three experimentally verified N-linked glycosylation sites at residues N57, N172, and N353. All three sites are utilized during protein maturation^61^. Using an estimated 693 F protein copies per virion, as determined from Equation (18), the total number of N-linked glycans associated with the F protein was calculated as described below:

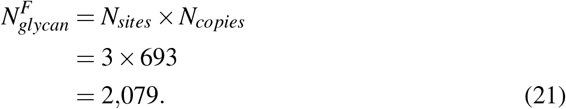

Therefore, the F protein provides 2,079 N-linked glycans to the virion.

#### 5.2.2 G Protein Glycosylation

The HMPV G protein is a mucin-like glycoprotein characterized by both N-linked and extensive O-linked glycosylation^24, 45^. The G protein copy number is estimated to be 139 per virion, based on calculation using Equation (19).

##### N-linked glycans

HMPV possesses between three and six N-linked glycosylation sites on its G protein^24^. For the soluble G (sG) protein of HMPV, five N-linked glycosylation sites are predicted^45^. Therefore, an average of four sites was used in this study to represent N-linked glycosylation. The number of N-linked glycans associated with the G protein was calculated as follows:

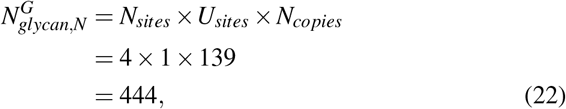

where *U*_*sites*_ = 1, as all N-linked glycosylation sites are utilized^61^.

##### O-linked glycans

The mucin-like ectodomain of the G protein contains multiple serine and threonine residues, which function as potential O-glycosylation sites^24^. Literature estimates suggest that the sG protein contains 59 O-linked sites^45^; however, not all sites are utilized, and the total number present on the G protein remains undetermined. Of these, 26 sites were identified as most frequently glycosylated on the G protein in HMPV ^24^. Consequently, 26 O-linked sites were included in this analysis. Utilization rates between 50 % to 100 % were evaluated; however, these variations resulted in only 1 % to 5 % changes in viral growth. Therefore, full utilization (*U*_*O*_ = 1) was assumed when calculating the number of O-linked glycans on the G protein in this study.

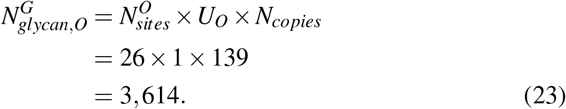

#### 5.2.3 Total Glycan Structures

Combining contributions from both glycoproteins (F, G) results in

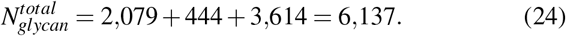

Therefore, a single HMPV virion is estimated to contain approximately 6,137 glycan structures.

#### 5.2.4 Monosaccharide Composition

Representative glycan structures were selected to calculate the total number of required monosaccharide units.

i. **N-linked glycans** were modeled as core-fucosylated, fully sialylated biantennary complex glycans, which represent a common mature N-glycan structure in mammalian systems and are denoted with the composition shown in Equation (25). This approach omits glycan microheterogeneity but provides a representative average composition for modeling.

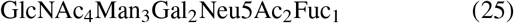 Equation (21) and Equation (22) are used to determine the total quantity of N-linked glycans present in the virion.

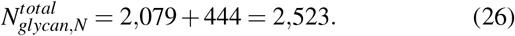 The total number of monosaccharide residues contributed by N-linked glycans is determined according to the following formula:

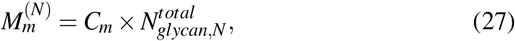

where *C*_*m*_ denotes the number of residues of monosaccharide *m* present in each glycan. For instance, the total number of N-linked galactose residues is calculated as follows:

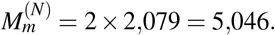
ii. **O-linked glycans** were modeled with the following composition.

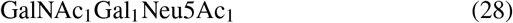 The total count of each O-linked glycan can be determined as follows.

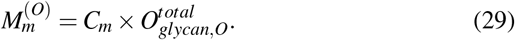 By substituting the values for galactose residues from Equation (28) and the total quantity of O-linked glycans from Equation (23) into Equation (29), we obtain 3,614 counts.
iii. **The final monosaccharide totals** were calculated by aggregating the contributions from both glycan types.

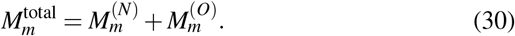 For instance, the total quantity of galactose residues was determined as follows:

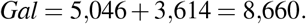 Table 13 presents a summary of the required monosaccharide counts, which are derived from N-linked and O-linked glycan structures in the HMPV virion.

### 5.3 Lipid Envelope Composition

Viruses acquire their envelope from host cell membranes during the budding process^62^, a mechanism that modifies the host cell lipid pathway^63^. Inhibition of specific lipids can suppress viral replication; for instance, sphingolipid inhibition reduced Human immunodeficiency viruse 1 (HIV-1) infectivity by a factor of four^62^. Therefore, viral lipid composition is a critical determinant in VBOF. Due to the scarcity of lipid composition data, lipids are often excluded from the Viral Biomass Objective Function (VBOF)^11^.

In this study, the lipid composition of the Sendai virus is utilized as a representative model for the paramyxovirus family, which includes HMPV. Analysis of the Sendai virus lipid composition revealed a phospholipid-to-cholesterol molar ratio of 1.0^19^, indicating equal quantities of cholesterol and phospholipids. Accordingly, the total lipid composition is defined as 50 % phospholipids and 50 % cholesterol. The phospholipid fraction consists of the following components: Phosphatidylcholine (PC) (37.7 %), Phosphatidylethanolamine (PE) (26.8 %), Phosphatidylserine (PS) (12.0 %), Sphingomyelin (SM) (8.8 %), Cardiolipin (CL) (9 %), Phosphatidylinositol (PI) (5.5 %), and Lysophosphatidylethanolamine (LPE) (1.6 %)^19^. The contribution of each phospholipid to the total lipid composition of the virus is presented in Table 14. Although the study on the Sendai virus used chicken eggs as the host, all lipids identified in the viral composition are also present in human cells ^20^. Therefore, the virus can also acquire its lipids from human cells.

**Table 14:**
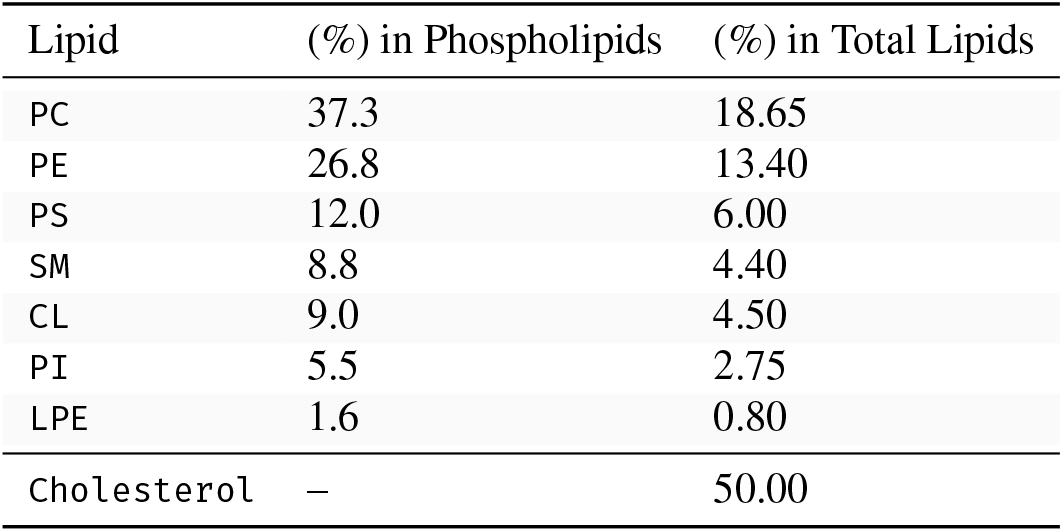
Lipid composition of the HMPV envelope. The phospholipid distribution was derived from the Sendai virus lipid analysis^19^. Phospholipids constitute 50 % of the total lipids, while cholesterol accounts for the remaining 50 %.

The total number of lipid molecules per virion was estimated based on the assumption of a representative spherical particle. A diameter of 200 nm^7^ and an average lipid packing density of 0.65 nm^2^ per lipid molecule^64^ in a bilayer were applied in the calculation, as outlined in the following formula:

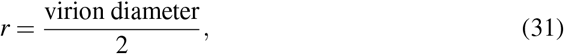

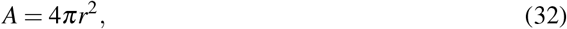

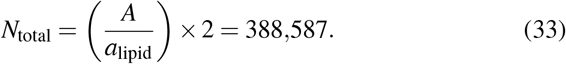

The lipid number was multiplied by 2 because HMPV possesses a lipid bilayer derived from the host cell membrane ^21^. This calculation resulted in a total of 388,587 lipid molecules per virion.

To estimate the number of molecules for each individual lipid species, their fractional abundance in the membrane was incorporated. The count of lipid species *i* is given by:

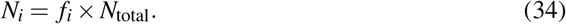

By substituting the total lipid count from Equation (33) into Equation (34):

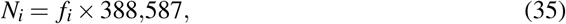

where, *f*_*i*_ represents the fraction of lipid species *i* within the total lipid composition. The lipid counts for individual lipid species are provided in Table 1.

### 5.4 VBOF Construction

The VBOF represents the stoichiometric requirements necessary for assembling a single unit of HMPV virion biomass and functions as the viral objective reaction within the integrated host-virus metabolic model. The VBOF was constructed using the methodology described by Aller et al.^11^ and was further extended to incorporate lipid and glycan components. The VBOF was developed by integrating five biochemical modules: (i) genomic RNA nucleotides, (ii) structural protein amino acids, (iii) bioenergetic costs of replication, (iv) envelope lipids, and (v) glycoprotein glycans.

#### 5.4.1 Genome Stoichiometry

The HMPV genome sequence (accession: NC_039199.1) and protein sequences with Reference Sequence (RefSeq)^65^ assembly accession, GCF_002815375.1 were obtained from the National Centre for Biotechnology Information (NCBI) database^66^.

Nucleotide composition was determined through direct sequence enumeration. The complete genome is 13,350 Kb in length. For spherical morphologies, one genome copy per virion was assumed, while two copies were assigned for filamentous particles, consistent with the packaging behavior reported in related pneumoviruses^37^.

#### 5.4.2 Protein Stoichiometry

Amino acid sequences for all nine HMPV structural proteins (N, P, M, F, M2-1, M2-2, SH, G, L) were retrieved from the NCBI RefSeq^65^ protein FASTA file GCF_002815375.1. Each protein sequence was analyzed to quantify the abundance of all 20 canonical amino acids. The stoichiometric requirement for each amino acid *j* was calculated as follows:

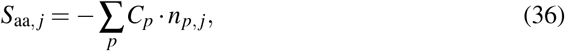

where *C*_*p*_ denotes the copy number of protein *p* per virion, as determined in Section 5.1, and *n*_*p, j*_ indicates the number of residues of type *j* in protein *p*. Negative coefficients indicate metabolite consumption.

#### 5.4.3 Bioenergetic Costs

The ATP and GTP requirements for virion production were determined based on standard biochemical reaction stoichiometries, shown in Table 15.

**Table 15:** Energy costs of individual biosynthetic processes.

| Process | Nucleotide | Cost per unit |
| --- | --- | --- |
| Genomic RNA synthesis | ATP | 2 per nucleotide |
| tRNA charging | ATP | 2 per amino acid |
| N-linked glycan attachment | ATP | 2 per N-glycan |
| O-linked glycan attachment | ATP | 1 per O-glycan |
| Ribosomal elongation | GTP | 2 per amino acid |
| Initiation and termination | GTP | 2 per protein |

Therefore, total energy requirements are as follows:

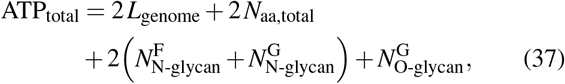

and

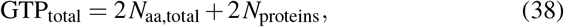

where *L*_genome_ denotes the genome length in nucleotides, *N*_aa,total_ represents the total amino acid count weighted by copy number, *N*_proteins_ indicates the number of distinct protein species (9 for HMPV), and *N*_N-glycan_ and *N*_O-glycan_ correspond to the total N- and O-linked glycan counts across all F and G protein copies, as described in Section 5.2.

The glycosylation ATP terms are thus directly linked to glycoprotein copy numbers and scale proportionally with virion morphology across variants. Hydrolysis byproducts (ADP, AMP, GDP, P_i_, H_2_O, H^+^) are also included as stoichiometric products, consistent with the VBOF framework of Aller et al.^11^.

#### 5.4.4 Lipids and Glycans stoichiometries

The stoichiometries of lipids and glycans are calculated in Section 5.3 and Section 5.2 VBOF, respectively.

#### 5.4.5 VBOF Assembly

The stoichiometries of the five components were consolidated into a single reaction by summing the coefficients of each metabolite across all modules. Cases in which metabolites appeared in both consumption and production terms were resolved through algebraic summation. The resulting reaction, designated as `HMPV_VBOF`, includes all metabolites with non-zero net coefficients and was exported in JavaScript Object Notation (JSON) format for subsequent normalization and integration into the host model (Section 5.5 and Section 5.6).

### 5.5 Normalization of VBOF Coefficients

The raw VBOF stoichiometric coefficients represent absolute molecular quantities. These coefficients are defined per virion and typically range from 10^3^ to 10^6^ molecules. However, these values are dimensionally incompatible with constraint-based modeling (CBM), where stoichiometric coefficients are conventionally expressed in millimoles per gram dry weight (mmol*/*(g_DW_ · h))^11, 12, 13, 14, 67^. Therefore, normalization to this unit basis is required. This normalization is essential for integrating the VBOF into a host cell genome-scale metabolic model and for ensuring that the resulting flux distributions, computed using FBA, remain physiologically interpretable.

The normalization procedure consists of two steps:

1. **Estimation of single-virion dry mass** - The dry mass of a single virion, *m*_v_ (g virion^−1^), is estimated from the raw stoichiometric coefficients as follows:

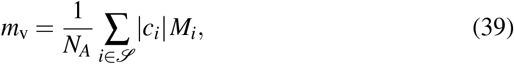

where *c*_*i*_ represents the raw coefficient, defined as the molecule count per virion for metabolite *i, M*_*i*_ is the molecular weight of metabolite *i* (g mol^−1^), and *N*_*A*_ = 6.022 *×* 10^23^ mol^−1^ is Avogadro’s number. *ℐ* denotes the set of all structural metabolites that constitute the virion. This set includes nucleotides, amino acids, lipids, and glycan precursors. All structural components contribute to *m*_v_, regardless of their stoichiometric sign in the reaction formulation. The use of the absolute value guarantees correct mass accounting.
2. **Conversion to** mmol gDW^−1^ - Each coefficient is subsequently recalculated on a per gram dry weight basis:

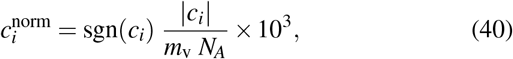

where 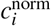 is expressed in units of mmol per gram of viral dry biomass (mmol gDW^−1^), consistent with biomass objective function conventions in GEMs. The function sgn(*c*_*i*_) preserves the directionality of the original coefficient, with negative values representing consumed substrates and positive values representing produced by-products. This formulation is equivalent to calculating the number of millimoles of metabolite *i* required to assemble one gram of virion particles. This approach aligns with the conventions of the biomass objective function established for GEMs in prokaryotic^67, 68^ and eukaryotic^67^ genomes.

### 5.6 Integration of VBOF into Host Model

The human bronchial epithelial cell metabolic model *i*HBEC^1^ was used as the host model, representing the primary target cell type for HMPV infection in the respiratory tract. This model comprises 2,047 reactions, 1,442 metabolites, and 1,390 genes. The model was imported using Constraints-Based Reconstruction and Analysis for Python (COBRApy)^69^. The model encompasses the full set of human metabolic reactions relevant to the respiratory epithelium, the primary site of HMPV infection, and incorporates a host biomass objective function (`biomass_hbec`) that specifies the stoichiometric requirements for host cell growth and maintenance.

#### 5.6.1 VBOF Reaction Construction and Addition

The resolved stoichiometry was used to instantiate the VBOF as a COBRApy `Reaction` object with the identifier `HMPV_VBOF`, assigned to the Viral Replication subsystem. This reaction was defined as irreversible, with bounds [0, 1,000], to reflect the unidirectional nature of virion assembly. The VBOF reaction was then appended to the model’s reaction list using the COBRApy `add_reactions` method. The resulting integrated model includes all host reactions as well as the single `HMPV_VBOF` reaction. This VBOF integrated model was exported in Systems Biology Markup Language (SBML) Level 3 format^70^ using the COBRApy^69^ `write_sbml_model` function for subsequent FBA^10, 71^ and flux variability analysis (FVA) analyses^72^.

### 5.7 Dual-Objective FBA for Antiviral Target Identification

Potential antiviral drug targets were identified through the application of a FBA framework to the integrated host HMPV metabolic model. Gene and reaction knockouts were conducted independently for each objective: one aimed to maximize the host biomass objective function (BOF) (`biomass_hbec`), and the other aimed to maximize the VBOF of HMPV (`HMPV_VBOF`). The effects of each knockout were subsequently evaluated by comparing outcomes across both objectives simultaneously. Prior to conducting the knockout screens, baseline fluxes for the wild-type (unmodified) condition were established. These baseline fluxes were determined for both objective functions by solving the model to optimality. Each objective function was optimized independently.

#### 5.7.1 Knockout Procedure

Each gene in the integrated model was individually knocked out using the COBRApy gene knockout function, which enforces gene-protein-reaction associations (GPR) rules. When a gene is deleted, the flux bounds of all reactions dependent exclusively on that gene are set to zero^69^. Each knockout was executed within a COBRApy context manager, which automatically reverts all changes after each simulation^69^. This approach prevents the accumulation of knockouts across iterations. For each gene *g*, knockout was implemented by setting its associated flux to zero. Two separate FBA were performed:

1. **Host growth assessment:** The BOF was maximized to determine the host growth rate following gene knockout.
2. **Virus replication assessment:** The VBOF was maximized to determine the viral output following gene knockout.

Results were expressed as percentages relative to the corresponding wild-type (unmodified) values. A value of 100 % indicates no measurable effect, while 0 % denotes complete loss of function. The following scale was applied:

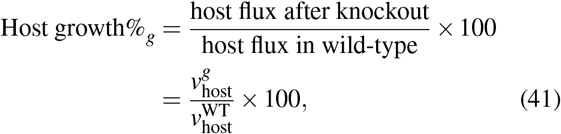

and

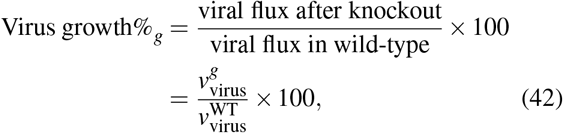

where 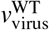 and 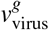 represent the viral growth rates before and after a specific gene knockout, respectively, while 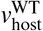 and 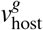 represent the host growth rates before and after the same gene knockout, respectively.

#### 5.7.2 Classifying the Impact of Each Knockout

Following the calculation of growth percentages for each knockout, each was classified into one of four impact categories according to the remaining flux relative to the wild-type (Table 16). A knockout with a score of 100 % indicated no measurable change, whereas a score of 0 % was considered lethal.

**Table 16:** Impact classification thresholds were applied for each knockout. These thresholds were applied independently to host and virus objectives.

| Category | Flux remaining | Interpretation |
| --- | --- | --- |
| lethal | < 1% | Gene/reaction is essential |
| significant | 1–50% | Strong impairment, but not fully lethal |
| moderate | 50–90% | Partial reduction in growth or replication |
| minimal | > 90% | knockout has little effect |

#### 5.7.3 Selectivity Scoring and Classification of Targets

Upon completion of both knockout processes, the results were combined using gene identifiers to generate a unified comparison table. Based on these results, the selectivity score was defined as follows:

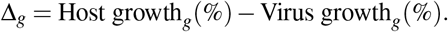

A large positive ∆_*g*_ value signifies effective host tolerance to the knockout, accompanied by a substantial impairment of viral replication. Genes were categorized into one of five mutually exclusive target classes based on the joint (host%, virus%) thresholds detailed in Table 17. The same procedure was applied to the reaction knockouts.

**Table 17:**
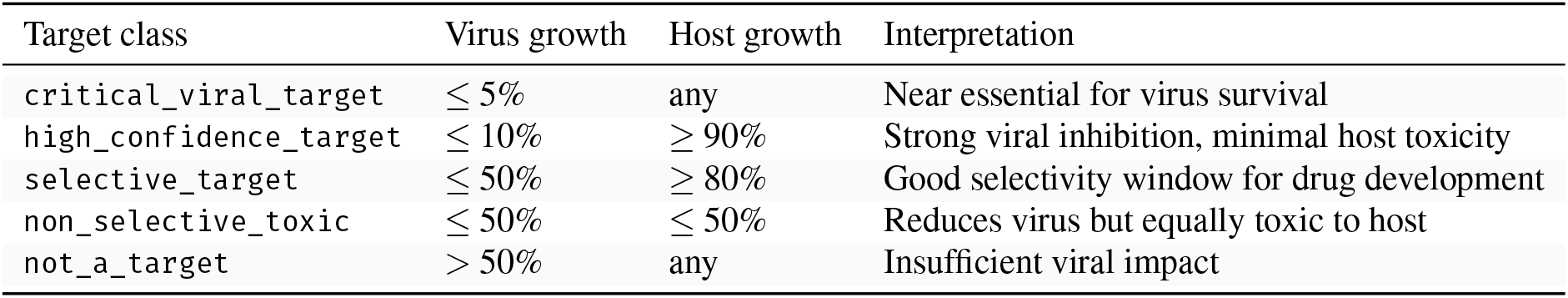
Antiviral target classification.

| Target class | Virus growth | Host growth | Interpretation |
| --- | --- | --- | --- |
| critical_viral_target | ≤ 5% | any | Near essential for virus survival |
| high_confidence_target | ≤ 10% | ≥ 90% | Strong viral inhibition, minimal host toxicity |
| selective_target | ≤ 50% | ≥ 80% | Good selectivity window for drug development |
| non_selective_toxic | ≤ 50% | ≤ 50% | Reduces virus but equally toxic to host |
| not_a_target | > 50% | any | Insufficient viral impact |

**Table 18:**
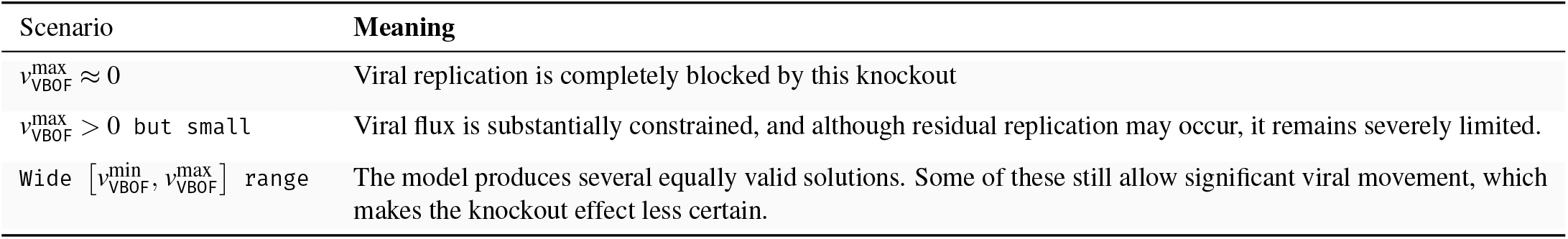
FVA on VBOF after knockout.

| Scenario | Meaning |
| --- | --- |
| $v_{VBOF}^{max} \approx 0$ | Viral replication is completely blocked by this knockout |
| $v_{VBOF}^{max} > 0$ but small | Viral flux is substantially constrained, and although residual replication may occur, it remains severely limited. |
| Wide $[v_{VBOF}^{min}, v_{VBOF}^{max}]$ range | The model produces several equally valid solutions. Some of these still allow significant viral movement, which makes the knockout effect less certain. |

### 5.8 FVA of Selective Antiviral Targets

Section 5.7 shows the potential antiviral targets by evaluating the single optimal FBA for each objective. However, a single optimal solution does not capture the full spectrum of possible outcomes^73^. Multiple flux distributions can achieve the same optimal value, and the biomass reaction may display substantial variation in flux ranges under these conditions^73^. To address this limitation, FVA was applied to each short-listed target^72^. FVA quantifies the minimum and maximum flux^73^ that the Viral Biomass Objective Function (VBOF) and the biomass objective function (BOF) can sustain following the knockout of a gene or reaction. This approach provides insight into the robustness of the predicted antiviral effect. Application of the FVA to each target produces four values: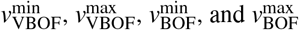

### 5.9 Software and Implementation

All source codes and the integrated model file are freely available from https://github.com/MSBIDynamics/HMPV-Host-VBOF. Table 19 presents the software and libraries used in this study, along with their respective versions.

**Table 19:** Software and libraries used for metabolic modeling and analysis.

| Software/Library | Version | Purpose |
| --- | --- | --- |
| Python<sup>74</sup> | 3.9.13 | Programming environment |
| COBRApy<sup>69</sup> | 0.26.0 | metabolic modeling and FBA |
| pandas<sup>75</sup> | 3.0 | Data processing and table manipulation |
| NumPy<sup>76</sup> | 2.4.2 | Numerical computations and operations |
| Matplotlib<sup>77</sup> | 3.8 | Data visualization and plotting |
| Seaborn<sup>78</sup> | 0.13.2 | Statistical data visualization |

## Data availability

The genome-scale metabolic model of a human bronchial epithelial cell, *i*HBEC^1^ infected with HMPV is available from BioModels Database^79^ under accession number MODEL2605050001 as a COMBINE archive OMEX file^80^ comprising an SBML Level L evel 3 Version 1 file^70^ with extension package flux balance constraints (fbc) version 2^81^. The Flux variability, Reaction deletion, Objective function, and Gene deletion (FROG) analysis framework was utilized to assess model reproducibility, ensuring reusability and results verification^9^.

## Code availability

Scripts to reproduce the analyses are available via GitHub in the repository https://github.com/MSBIDynamics/HMPV-Host-VBOF (Zenodo-Archive: 10.5281/zenodo.20628356 under the terms of the Massachusetts Institute of Technology (MIT) license from.

## Author contributions

R.M. and B.S. developed the conceptual idea. R.M. derived the computational method. S.H. implemented the method and performed the analysis. S.H. created the model and analyzed it. S.H. created the GitHub repository to follow the FAIR (Findability, Accessibility, Interoperability, and Reusability) principles. S.H. designed all figures and tables. R.M. supervised the work and wrapped the model in an Open Modeling EXchange format (OMEX) archive, implemented FROG analysis on the model, and uploaded it to BioModels. S.H. and L.B. wrote the manuscript. R.M. and B.S. critically revised the manuscript and the figures. All authors have revised, read, and accepted the manuscript in its final form.

## Acknowledgments

This work was funded by the *Forschungscampus Mittelhessen* (FCMH, Research Campus of Central Hessen). The authors acknowledge the support by the Open Access Publishing Fund of Justus Liebig University Giessen (https://www.uni-giessen.de/ub/en/publish/openaccess-en).

Bernd Schmeck reports funding from the German Center for Lung Research (DZL 4.0), from Bundesministerium für Forschung, Technologie und Raumfahrt (Federal Ministry of Research, Technology and Space; BMFTR) for CALM-QE within the Medical Informatics Funding Scheme (FKZ 01ZZ2318A), PermedCOPD (FKZ 01EK2203A), and the Hessisches Ministerium für Wissenschaft und Forschung, Kunst und Kultur (LOEWE Habitat and LOEWE Diffusible Signals).

## List of Abbreviations

AA: angstrom
ACGAMPM: Phosphoacetylglucosamine mutase
ACGAM6PS: GlcNAc-6-P synthase
AHCi: Adenosylhomocysteinase
ATP: adenosine triphosphate
ADP: adenosine diphosphate
AMP: adenosine monophosphate
BiGG: Biochemical, Genetical, and Genomical
BOF: biomass objective function
CBM: constraint-based modeling
CL: Cardiolipin
COBRApy: Constraints-Based Reconstruction and Analysis for Python
COMBINE: Computational Modeling in Biology Network
COVID-19: Coronavirus Disease 2019
cryoEM: cryo–electron microscopy
cryo-ET: cryogenic electron tomography
DON: 6-Diazo-5-oxo-L-norleucine
F: fusion protein
FCMH: Forschungscampus Mittelhessen
FBA: flux balance analysis
fbc: flux balance constraints
FROG: Flux variability, Reaction deletion, Objective function, and Gene deletion
FVA: flux variability analysis
G: glycoprotein
GEM: genome-scale metabolic model
GFPT: Glutamine—fructose-6-phosphate aminotransferase
GF6PTA: Glutamine-fructose-6-phosphate transaminase
GK1: Guanylate kinase 1
GPR: gene-protein-reaction associations
GTP: Guanosine Triphosphate
GDP: Guanosine diphosphate
GMP: Guanosine Monophosphate
HC: High Confidence Target
HIV: Human immunodeficiency viruses
HIV-1: Human immunodeficiency viruse 1
HBP: hexosamine biosynthesis pathway
HMPV: Human Metapneumovirus
hRSV: Human respiratory syncytial virus
ID: identifier
JSON: JavaScript Object Notation
Kb: kilobase
L: large polymerase protein
LPE: Lysophosphatidylethanolamine
M: matrix protein
METAT: Methionine adenosyltransferase
MIT: Massachusetts Institute of Technology
mRNA: messenger ribonucleic acid
N: nucleoprotein
NCBI: National Centre for Biotechnology Information
NT: nucleotide metabolism nm nanometer
OMEX: Open Modeling EXchange format
RCSB: Research Collaboratory for Structural Bioinformatics
P: phosphoprotein
PC: Phosphatidylcholine
PE: Phosphatidylethanolamine
PI: Phosphatidylinositol
PS: Phosphatidylserine
PDB: Protein Data Bank
RefSeq: Reference Sequence
RNA: ribonucleic acid
RdRp: RNA-dependent RNA polymerase
RNP: Ribonucleoprotein
RSV: Respiratory Syncytial Virus
S: Selective Target
SAM: S-adenosylmethionine
SARS-CoV-2: Severe Acute Respiratory Syndrome Coronavirus 2
SBML: Systems Biology Markup Language
siRNA: Small interfering RNA
sG: soluble G
SH: small hydrophobic protein
SM: Sphingomyelin
UAGDP: UDP-GlcNAc diphosphorylase
UDP: Uridine Diphosphate
VBOF: Viral Biomass Objective Function
WHO: World Health Organization

